# Novel genes arise from genomic deletions across the bacterial tree of life

**DOI:** 10.64898/2026.01.05.697752

**Authors:** Arya Kaul, Fernando Rossine, Karel Břinda, Michael Baym

## Abstract

Bacteria are hosts to enormous genic diversity. How new genes emerge, functionalize, and spread remain longstanding questions. Here, we explore a mechanism by which adaptive deletions fuse distant gene fragments. Unlike other gene birth mechanisms that begin with rare, neutral mutations, these “deletion-born fusions” reach high frequency by hitch-hiking on the deletion. The deletion-driven proliferation of the fusion prolongs the mutational supply within these genes before loss, providing additional opportunities for neofunctionalization. We document one such gene fixing and expressing in a long-term *E. coli* evolution experiment, and identify additional fusion events in the *Mycobacterium tuberculosis-bovis* split. Finally, we develop a scalable systematic screen to detect these genes in all 2.4 million public single-isolate genomes and identify deletion-born fusions across the bacterial tree of life. These findings challenge the notion that deletions are solely destructive and highlight their role as potential catalysts for evolutionary innovation.

## INTRODUCTION

Bacteria are the most genetically diverse domain of life on Earth.^1,2^ This genetic diversity also translates to a profound diversity in their protein-coding genic repertoire. New genes and gene families continually arise across the bacterial tree of life; pangenome studies of single species can identify tens of thousands of protein families, and metagenomic sequencing of environments like the human gut has revealed millions of protein-coding genes of unknown function.^3–7^ This diversity raises the question of where new bacterial genes and their corresponding functions come from.

Different mechanisms for bacterial gene birth have been proposed and each uniquely constrains the generation of functional novelty. On the one hand, duplication and subsequent diversification ensures that novel genes derive from functional templates,^8–10^ but the preexisting functionality and domain structure constrain their evolutionary potential. On the other hand, overprinting,^11,12^ the expression of a novel gene overlapping a previously existing sequence but in a different reading frame, generates protein products without homology to existing proteins or prior functional constraints. Yet new proteins created from random peptides often suffer from excessive hydrophobicity, aggregation propensity, and intrinsic disorder.^13,14^ Considering these tradeoffs, gene fusion, the stitching together of entire fragments of distinct genes, provides a gene birth mechanism that balances innovation with functionality by bringing together existing protein domains into new contexts,^15^ though such fusions must still avoid being lost to selection or genetic drift.

Random loss of newborn genes is compounded a deletional bias that threatens to purge them from bacterial genomes before they can acquire beneficial function. Compact bacterial genomes, often less than fifteen percent non-coding,^16^ reflect a pervasive bias favoring loss of non-essential DNA.^17–19^ This process, seen in natural,^20^ clinical,^21,22^ and experimental settings,^23,24^ is typically framed as a loss of genetic potential. These deletions may arise through diverse mechanisms, including homologous recombination,^25^ replication-associated slippage,^26,27^ erroneous repair,^28^ or site-specific recombinases.^29^ Regardless of mechanism, deletions are a hallmark of bacterial genome evolution and can be positively selected by reducing the metabolic cost of replicating DNA or by tuning gene interaction networks.^16,19,30^

Deletions can also generate new arrangements of existing material. In a recent analysis of the Lenski Long-Term Evolution Experiment (LTEE),^31^ uz-Zaman et al. found that deletions moving regulatory elements contributed the largest share to the transcription and translation of previously non-coding DNA.^32^ In other instances, deletions generated functional gene fusions that led to new anti-phage defense functions,^33^ to phenotypes with increased colony spread,^34^ and to potentially adaptive gene products in *Mycobacterium tuberculosis* lineages.^35^ Despite the importance of gene fusions in generating novel phenotypes and individual cases linking deletions to the emergence of fusion genes, it remains unknown whether this represents a widespread mechanism for bacterial genome innovation.

Here, we explore a model of bacterial gene birth in which a deletion results in the fusion of the start of one open reading frame (ORF) and the end of another to create a novel ORF. Next, we identify the creation and maintenance of these deletion-born fusions in two densely sampled contexts: in the Lenski LTEE, and during mycobacterial speciation. Finally, we develop a computational technique to efficiently query putative deletion-born fusions across multi-million bacterial isolate genome collections and find evidence for this mechanism of gene birth across the bacterial tree of life. Our findings reframe the bacterial deletional bias as not merely destructive, but as a possible creative force.

## RESULTS

### Fusion genes can spread by hitchhiking on the fitness benefit of their causal deletion

Deletions in a bacterial genome result in the fusing of two distal sections of genetic material spanning the deletion junction. Either one or both deletion boundaries can fall inside an existing gene, thus generating a chimeric ORF at the junction. This is made more likely by the gene-dense architecture of bacterial genomes.^36^ Both the novel ORF and the removal of intervening material can cause changes in relative fitness.

If a deletion confers a fitness benefit, any fusion ORF created as a by-product can increase in frequency through hitchhiking. Although most nascent genes are expected to be neutral or deleterious regardless of the mechanism by which they originated,^37^ deletion-born fusions may possess an advantage over other novel genes. Unlike mechanisms that depend on rare, initially neutral genes gradually drifting to appreciable frequency, deletion-born fusions can be quickly driven to elevated frequencies as direct correlates of positively selected structural changes. This hitchhiking may prolong their residence time in the population, increasing the likelihood that subsequent mutations will convert them into beneficial, functional alleles before being lost to drift or purifying selection (**Figure 1**). Moreover, because these fusions are assembled from pre-existing coding material, they are more likely to fold into active catalytic, structural, or regulatory domains. Overall, we expect that deletion-born fusion genes should be more likely to be functionalized than novel genes that emerge through different mechanisms.

**Figure 1.**
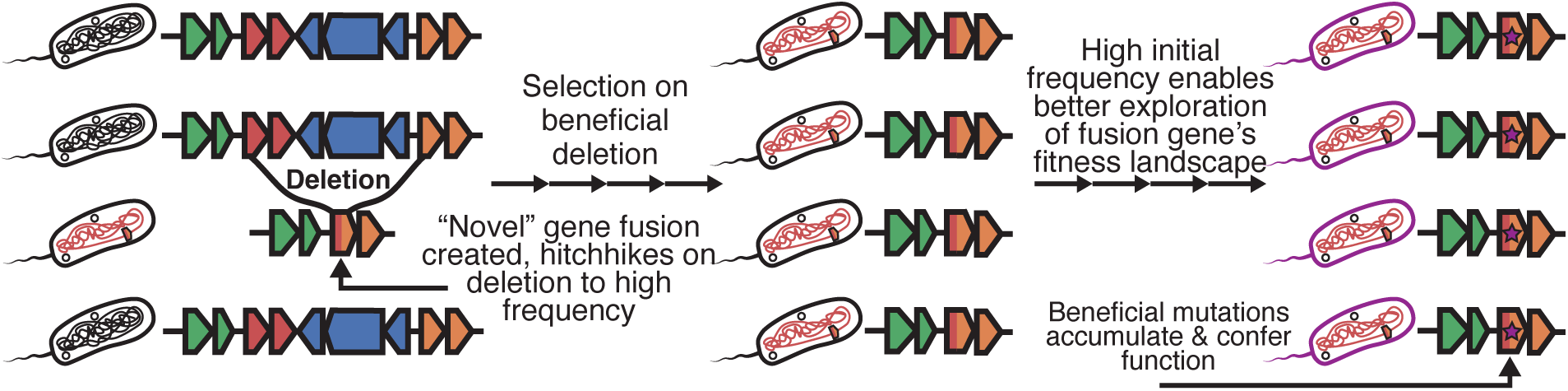
Proposed model of deletion-born fusion genes. (1) An initial deletion occurs in a subset of the population and spontaneously creates a deletion-born fusion gene, (2) The deletion is beneficial and rises in frequency with the fusion gene hitchhiking to high frequency, (3) fusion gene at high initial frequency explores fitness landscape and functionalizes

To formalize this verbal model, we constructed a stochastic simulation that incorporates the role of genetic hitchhiking and allowed us to contrast the functionalization likelihood of novel genes that arose through different mechanisms. We found that starting novel gene frequency is the dominant driver of neofunctionalization, supporting the hypothesis that hitchhiking on a deletion is a means by which novel ORFs can functionalize (**Supplementary Figure 1**). We constructed a forward-time Wright-Fisher simulation with 10^6^ members, each with one of two states (no novel gene, and non-functional novel gene present with cost *c*). We assumed that each novel gene is initially non-functional, could be purged each generation with probability *p_purge_*, and could functionalize with a probability *µ*. We also assumed that deletion-born fusion genes quickly increased in frequency to some frequency p*_init_* (which should be reflective of the fitness advantage of the genomic deletion that gave rise to the gene) while genes born from other mechanisms always had *p_init_* = 10^-6^ (a lone member of the population begins with the gene). We demonstrated that across values for *p_purge_* and *µ* the greatest determinant of at least one member of the population functionalizing the gene before it is purged from the population is *p_init_*, the initial frequency of the gene. When *p_init_* = 10^-6^ the gene never functionalized consistently; however, as *p_init_* increased, the probability of functionalizing increased substantially. This functionalization probability still fell below many values of *µ*; especially if the fusion is deleterious instead of neutral. It was only when *p_init_* approached 1 that neofunctionalization occurred consistently under a sweep of parameters. These results suggested that young deletion-born fusion genes should be present in populations that are adapting to new environments, when genomic deletions are typically the most advantageous (i.e., *p_init_* is the highest).

### A novel deletion-born fusion fixes in the Lenski LTEE

To investigate the emergence of deletion-born fusions, we analyzed data from the Lenski LTEE. Briefly, the LTEE tracks the evolution of 12 replicate populations of *Escherichia coli* in minimal media since 1988, spanning over 82,000 generations as of 2025.^38,39^ By clustering predicted ORFs from sequenced clonal isolates against the ancestral genome, we identified a novel fusion gene created by a 57.5 kb deletion in the Ara+1 lineage. This deletion fused *yjcO* (a gene of unknown function) to *lysU* (lysyl-tRNA synthetase) (**Figure 2A**).

**Figure 2.**
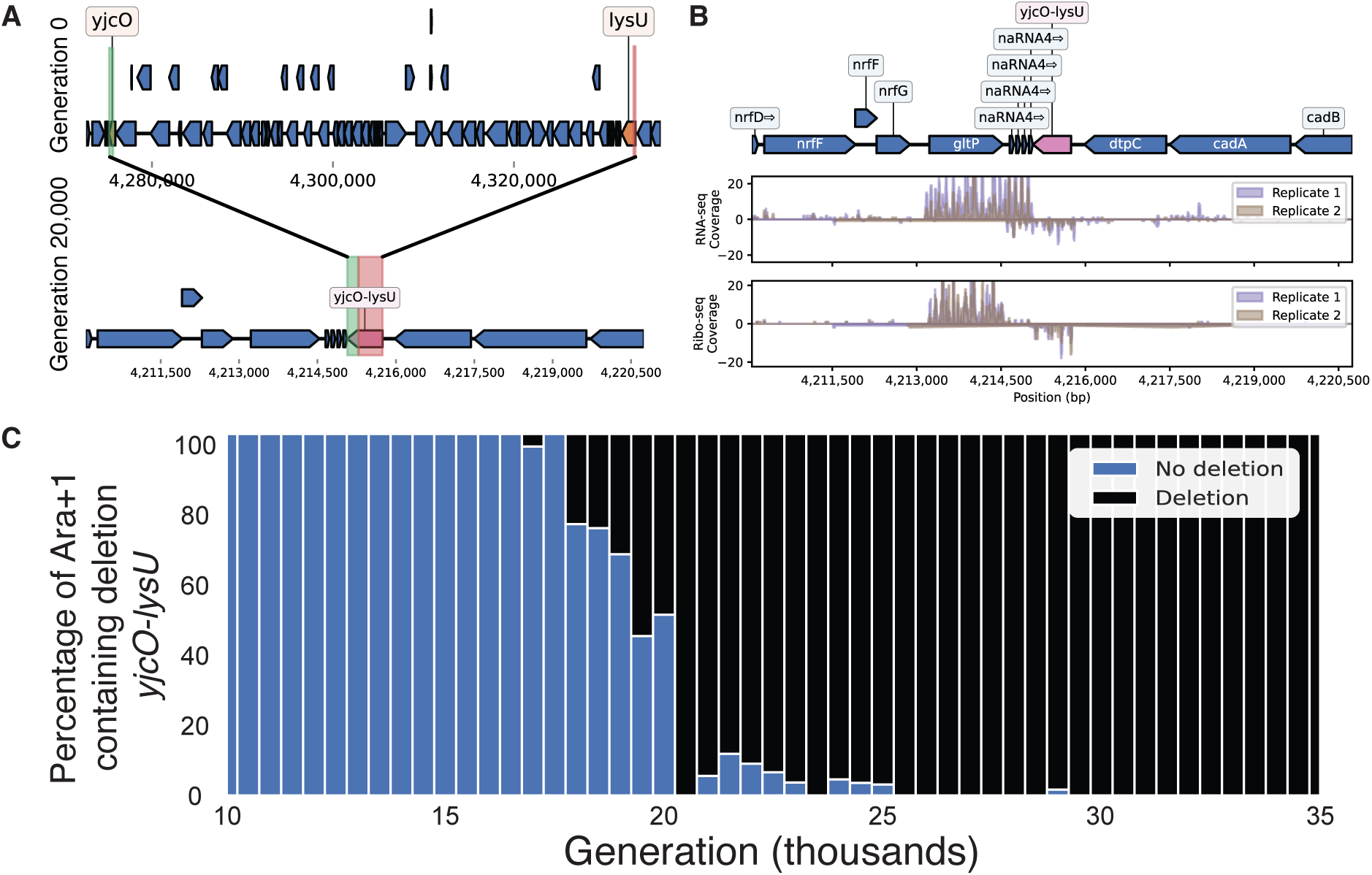
A 57.5 kb deletion creates a fusion gene in the Lenski LTEE Ara+1 lineage. (A) Top: Genomic locus surrounding *yjcO-lysU* in REL606, Bottom: The same genomic locus of the clonal isolate from Generation 20,000 with the deletion and resulting *yjcO-lysU* fusion gene. (B) Genomic locus of the *yjcO-lysU* region with coverage plots corresponding to RNA-seq and Ribo-profiling data from two clonal isolates picked at generation 50,000 in Ara+1. (C) Metagenomic sequencing results comparing the fraction of reads supporting the deletion or not between Generation 10,000 and generation 35,000 in the Ara+1 lineage.

The fusion of *yjcO* and *lysU* is expressed and potentially folds but is likely non-enzymatic and functionally inert. Re-analysis of existing RNA-seq and ribosome profiling data from clonal isolates in Ara+1 revealed that the *yjcO-lysU* fusion is both expressed and translated at generation 50,000 (**Figure 2B**).^40^ Domain annotation shows that the fusion retained multiple Sel1-like repeats from *yjcO*, but no catalytic domains were preserved from *lysU*. Across all clonal isolates sequenced after its appearance, the fusion gene shows no nucleotide substitutions. Based on the gene length and known LTEE mutation rates, we estimate the probability of at least one mutation occurring after its appearance to be <1%, consistent with the observed absence of variation and suggesting weak or no selection on the fusion.

Both the fusion and the deletion it arose from are first detected in metagenomic sequencing data at around generation 19,500 and appear to fix within ∼500 generations (**Figure 2C**). The speed of this selective sweep indicates that either the deletion itself or the resulting *yjcO-lysU* fusion likely conferred an approximate 6.5% fitness advantage under LTEE conditions (**Supplementary Note**).

Supporting the hypothesis that the selective advantage lies in the deletion itself rather than the fusion gene, a parallel 43.4 kb deletion independently arose and fixed in the Ara-3 lineage at the same genomic locus, but did not produce a novel fusion (**Supplementary Figure 2**).

Additional analyses of transposon sequencing data confirmed that disrupting the fusion gene had no deleterious effect,^41^ reinforcing the interpretation that the deletion, not the gene, is the beneficial variant.

Genomic regions flanking deletions ≥1 kb exhibited significantly greater and more variable changes in expression and translation than randomly sampled regions (**Supplementary Figure 3**). These effects decayed with increasing window size, consistent with deletions inducing localized promoter capture and the disruption of transcriptional context. This supports a model in which deletions remodel local expression and translational landscapes, occasionally generating novel transcribed or translated products.

These results indicate that, even in the relatively constant conditions of the LTEE, new fusion genes can arise from large deletions and quickly increase in frequency in the population.

### Deletion-born fusions emerge during speciation in the Mycobacterium Tuberculosis Complex

We next turned to a well-characterized and deeply sequenced bacterial speciation event, the *Mycobacterium tuberculosis*-*bovis* divergence, to investigate the potential for deletion-born fusions to arise in complex natural systems.^42,43^

We identified two putative deletion-born fusion genes: *acrR-glcD* (from a 2 kb deletion) and *mlaE-htpX* (from a 12.5 kb deletion) arising during this divergence (**Figure 3**). Using a collection of 47 long-read assemblies (37 *M. tb*, 10 *M. bovis*), we predicted all ORFs, clustered them into orthologous groups, and searched for novel ORFs consistent with deletions (see METHODS).

**Figure 3.**
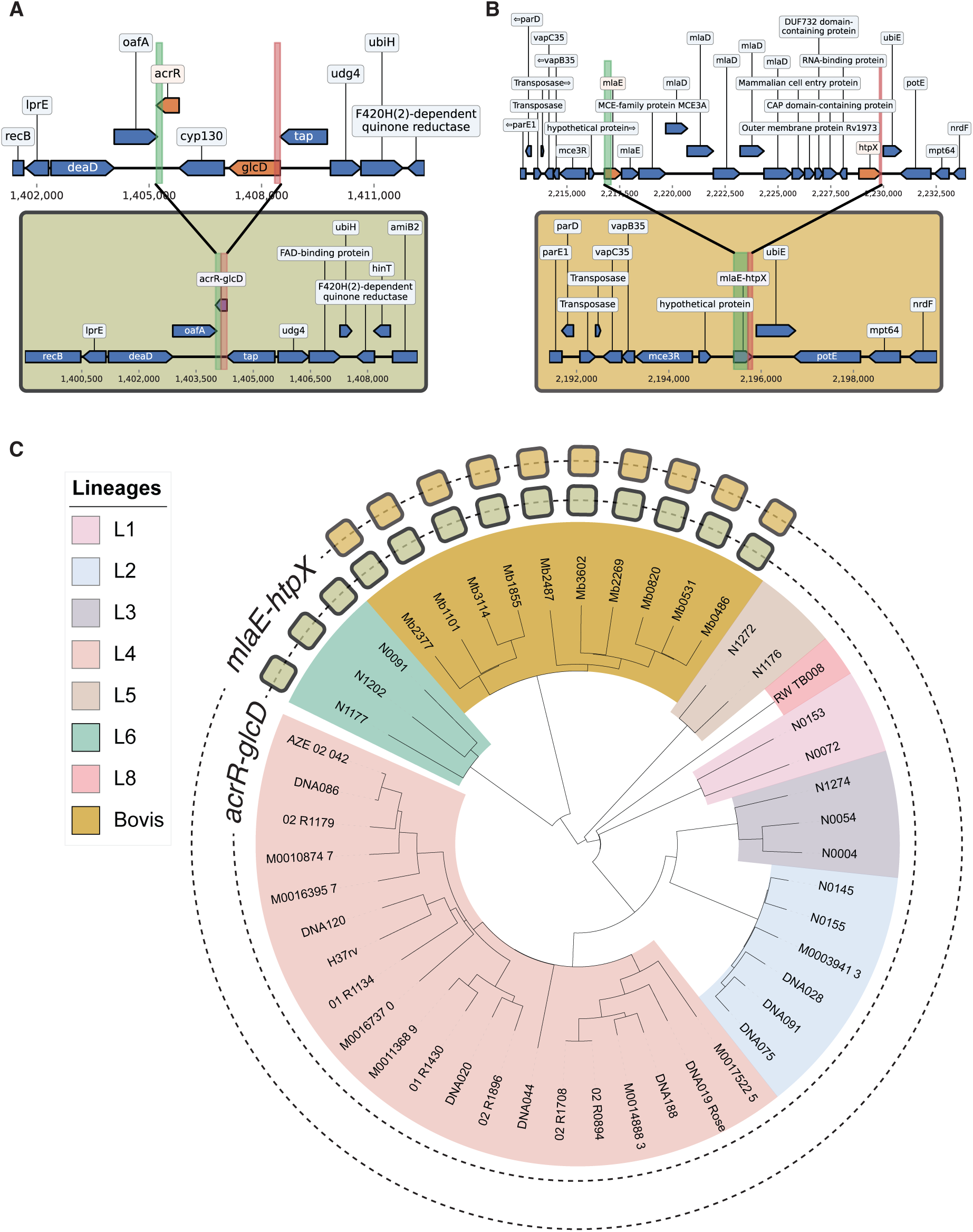
Two deletion-born fusions arose in the M. tb/M. bovis speciation event. (A) Top: representative pre-deletion *acrR-glcD* genomic neighborhood in *M. tb*. (genome: RW-TB008). Bottom: representative post-deletion *acrR-glcD* genomic neighborhood in *M. bovis* + L6 *M. tb*. (genome: Mb1855). (B) Top: pre-deletion *mlaE-htpX* genomic neighborhood in *M. tb*. (genome: DNA019_Rose). Bottom: post-deletion *mlaE-htpX* genomic neighborhood in *M. bovis* (genome: Mb3602). (C) Phylogenetic tree of the 47 analyzed samples. Clade colors denote the lineage each isolate falls within, and the filled squares in each ring denote the genomes with *acrR-glcD* and *mlaE-htpX* respectively.

Phylogenetic mapping showed that *mlaE-htpX* occurred exclusively in *M. bovis*, while *acrR-glcD* was found in both *M. bovis* and *M. tb* Lineage 6, the closest relative of *M. bovis*.^44^ These monophyletic distributions suggest each fusion arose once, *mlaE-htpX* in the *M. bovis* split and *acrR-glcD* in the *M. bovis*/L6 divergence.

Neither fusion appears to be immediately functional. Structurally, *acrR-glcD* maintains reading frame continuity between the N-terminus of *glcD* and the C-terminus of *acrR*, whereas *mlaE-htpX* fuses *mlaE* in-frame to an out-of-frame fragment of *htpX*. Pfam domain searches revealed that both fusions lost the catalytic motifs of their ancestors and did not generate any new conserved domains. While neither *acrR-glcD* nor *mlaE-htpX* exhibit nucleotide variation in the genomes from Marin et al. and Charles et al.,^42,43^ a broader analysis (next section) revealed both variation and signatures of selection.

The observation of deletion-born fusions in the twin contexts of laboratory evolution and natural speciation in evolutionarily distant bacteria implies that this process may be widespread across bacterial diversity.

### An alignment-free approach to characterize structural variation across a multi-million bacterial genome collection

To query structural variation at the scale of the bacterial tree of life, our previous alignment-based techniques were computationally infeasible. We therefore developed an alignment-free approach that queries short k-mers from the beginning (“prefix”) and the ending (“suffix”) of candidate genes to infer structural rearrangements based upon the genomic distance between these sequences (**Figure 4A**). Because this method relies only on the location of exact k-mer matches, it can leverage FM-indices for sublinear query times,^45^ making searches across millions of genomes tractable. We applied this approach to AllTheBacteria (ATB), the largest current bacterial genome collection, comprising 2.4 million uniformly assembled single isolate genomes.

**Figure 4.**
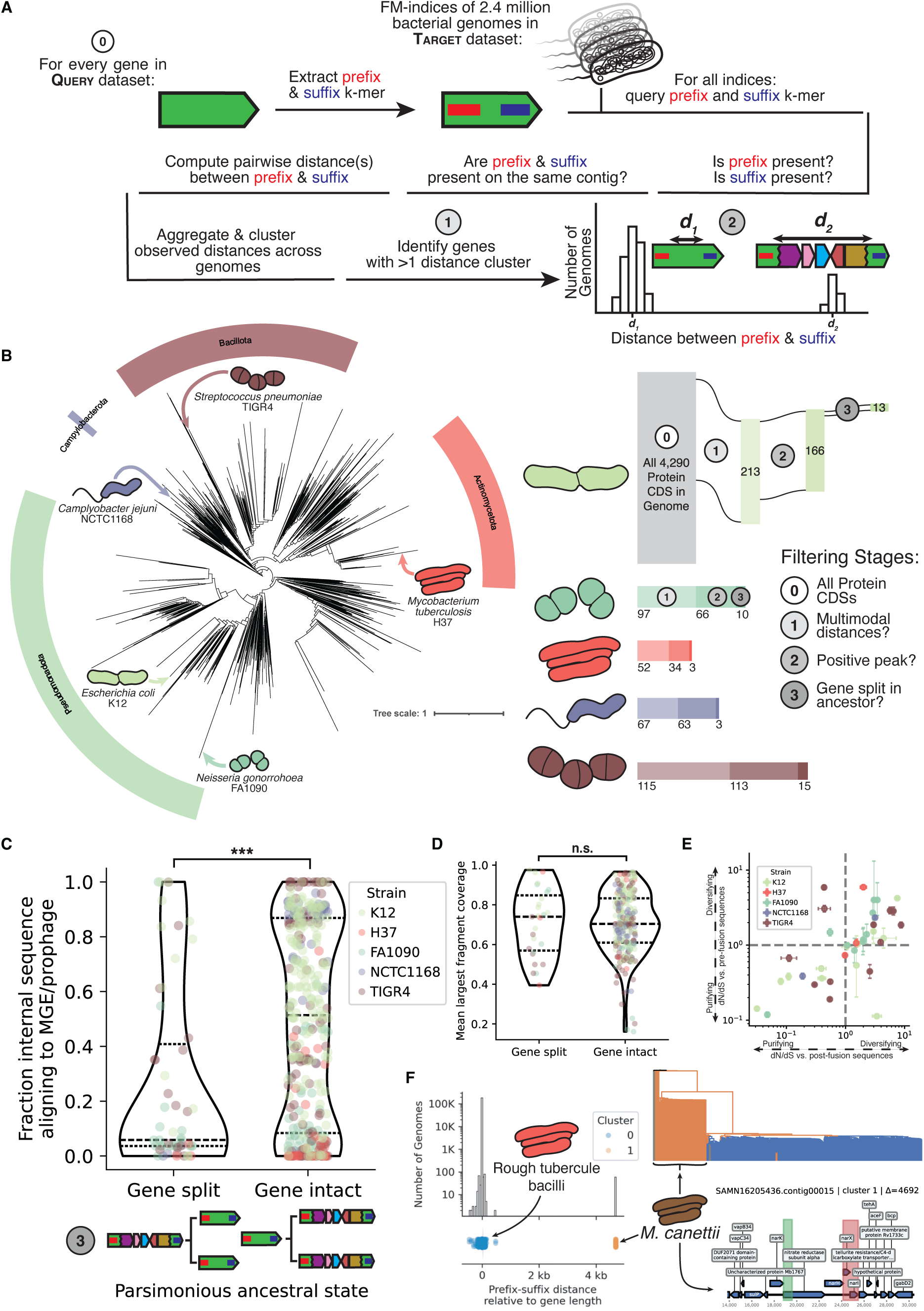
Deletion-born fusions are present across the bacterial tree of life. (A) Schematic of the prefix-suffix k-mer approach. (B) Left: bacterial tree of life condensed to the genus level showing the five type strains queried. Right: filtering cascade with counts of genes passing each stage: (0) all protein-coding ORFs, (1) multimodal prefix-suffix distances, (2) positive distance peak present, (3) gene inferred as split in ancestor. Bacterial illustrations were traced from SEM images. *E. coli*,^46^ *N. gonorrhoeae*,^47^ *M. tuberculosis*,^48^ *C. jejuni*,^49^ *S. pneumoniae*.^50^ (C) Fraction of intervening sequence aligning to MGEs/prophages in genes with split versus intact ancestral states (Mann-Whitney U test, p < 10⁻⁵). (D) Mean coverage of the largest alignment block between prefix and suffix k-mers; no significant difference between distributions. (E) dN/dS estimates for 44 candidate fusions. X-axis: dN/dS versus intact alleles; Y-axis: dN/dS versus reconstructed pre-deletion sequences. Error bars show 95% confidence intervals. (F) *narX* from *M. tuberculosis* H37Rv. Histogram and stripplot show prefix-suffix distances; Cluster 0 = rough tubercle bacilli (including the fusion), Cluster 1 = *M. canettii* (putative ancestral split state). Phylogeny colored by cluster assignment; bottom panel shows the 10 kb genomic neighborhood of the ancestral *narX* locus in *M. canettii*.

The distribution of observed prefix-suffix distances reflects structural variation within a given gene. Nearly all insertion, deletion, or partial inversion events will change the distribution of distances. For deletions, the ancestral pre-deletion sequence will have a larger prefix-suffix distance than a fused post-deletion sequence. We used multi-modality in the prefix-suffix distance distribution as an indicator of structural variation within a locus (see METHODS).

We first validated this prefix-suffix approach on the three previously identified fusion genes and observed distinct peaks at the ancestral and fused distances in all cases (**Supplementary Figure 4**). Notably, the scale of ATB revealed additional structural variation at these loci that was invisible in our earlier, smaller-scale analyses. At the *yjcO-lysU* locus, we identified nine distinct distance clusters spanning a range of structural variants. The 80 genomes in cluster 0 (prefix-suffix distance matching the fusion length) formed a monophyletic clade restricted to LTEE-derived isolates. The remaining 116,892 genomes showed extended prefix-suffix distances ranging from 29 to 96 kb, consistent with independent insertions and deletions at this locus. Cluster 3, comprising 48,892 genomes with a prefix-suffix distance centered at 57 kb, matches the ancestral LTEE length; the remaining 68,000 genomes exhibit distinct structural configurations, revealing additional diversity at this locus (**Supplementary Figure 4A**).

In *Mycobacteria*, the deeper sampling from ATB uncovered 4 distinct *acrR-glcD* alleles and 17 *mlaE*-*htpX* alleles. Alignment-based Bayesian selection analysis showed strong evidence for diversifying selection at codon 50 in *acrR-glcD* (posterior >0.9) and purifying selection at five positions (87, 91, 92, 108, 125) in *mlaE-htpX* (see METHODS). These patterns indicate that, even absent recognizable catalytic domains, these fusions are experiencing selective pressure, consistent with emerging or maintained function.

While we developed the prefix-suffix method to identify deletion-born fusions, the approach detects any recurrent structural variation at a locus. In the next section we filter candidates to deletion-born fusions, but many of the events we filter out are themselves biologically interesting. We note the identification of internal deletions in the *mngB* gene in *E. coli* (**Supplementary Figure 5A**), repeat prophage insertions into *rep13e12* gene in *M. tuberculosis* (**Supplementary Figure 5B**), and variable gene cargo in an uncharacterized mobile element disrupting the *pdp* gene in *S. pneumoniae* (**Supplementary Figure 5C**). Thus, the prefix-suffix signal is generalizable to surveying structural variation beyond our specific application and, because of its computational efficiency, scales readily to ever-expanding genome collections.

### Putative deletion-born fusions are found across the bacterial tree of life

We next asked how pervasive deletion-born fusions are across diverse bacterial phyla. We applied the prefix-suffix k-mer screen to all annotated protein-coding sequences from five type strains spanning well-studied and diverse bacterial clades (*Escherichia coli* K12, *Mycobacterium tuberculosis* H37Rv, *Neisseria gonorrhoeae* FA1090, *Campylobacter jejuni* NCTC1168, and *Streptococcus pneumoniae* TIGR4). We selected these species since they span phylogenetically distinct clades, and because their clinical importance has made them among the most deeply sequenced bacteria, with each represented by at least 60,000 genomes in the ATB.

Across the strains, our pipeline resolved 44 putative deletion-born fusion candidates inferred as “split” at the MRCA of the genomes sampled: 13 in *E. coli* K12, 3 in *M. tuberculosis* H37Rv, 10 in *N. gonorrhoeae* FA1090, 3 in *C. jejuni* NCTC1168, and 15 in *S. pneumoniae* TIGR4 (**Figure 4B**, **Supplementary Table 1**).

We identified these putative deletion-born fusions via a sequence of consecutive filters: We first selected genes whose prefix-suffix distances were multimodal, indicative of structural variation. We then retained candidates that showed one cluster centered near the length of the gene (a difference of 0 implying an intact gene) and at least one more cluster at a larger, positive distance (putative pre-deletion sequence) (see METHODS). By solely relying on the distance between the prefix and suffix k-mers, we are unable to distinguish between insertion into a gene and a deletion that led to the formation of that gene. To distinguish between these possibilities, for each candidate gene, we extracted complete genomes with equal representation from each cluster, added outgroups, built a k-mer–based distance tree, and performed parsimony-based ancestral state reconstruction using cluster labels as discrete states (see METHODS). Candidates were retained when the most parsimonious cluster assignment for the most recent common ancestor (MRCA) of all sampled genomes was a cluster with a positive distance length (split gene) implying that the ancestral state was an unfused gene that later experienced a deletion and gene formation.

In genes whose parsimonious ancestral state is predicted to be intact, the observed positive prefix-suffix distance peaks are significantly explained by the insertion of foreign elements, particularly mobile genetic elements (MGEs) and prophages (**Figure 4C**). By contrast, at loci whose MRCA is inferred not to have carried the intact gene, the intervening segments have significantly fewer matches to MGEs or prophages, consistent with separation/fusion via deletion or recombination rather than insertion.

To ensure the distance signals identified were not derived from spurious k-mer matches, we sequence-aligned the putative fusion gene to genomes where it was predicted to be split. We found that all putative deletion-born fusions have at least 80% gap-excluded identity to their split ancestors, implying the prefix-suffix approach detected true sequence homology. Further, the distribution of the relative contributions of the largest alignment fragment in putative deletion-born fusions (MRCA = Gene split) matched that of disrupted genes (MRCA = Gene intact, which we expect to be fully random), implying that spurious k-mer matches to either the prefix or suffix alone are undetectably rare (**Figure 4D**).

The lack of spurious k-mer matches is likely attributed to the significant requirements we enforce: at least 54 base pairs of exact nucleotides matching on the same contig separated by roughly the same amount across numerous genomes. To test how robust this approach was to varying lengths of k we also tested numerous values for three sets of 1,000 random bacterial RefSeq proteins. We found that below k = 20, the number of genes with multimodal distributions rises sharply (**Supplementary Figure 6**). Though some of these signals might be real, we elected to continue with the more stringent value of k = 27 to ensure high confidence matches.

Signatures of positive selection are most common across the 44 putative deletion-born fusions (**Figure 4E**). To distinguish between selection currently operating on the proteins and strong selection immediately after the initial deletion event, we categorized synonymous and nonsynonymous variation in two ways. First, we sampled genomes with the fusion gene and measured their variation against the reference fusion gene (dN/dS vs. post-fusion sequences). Second, we measured variation against “surrogate” genes created by merging the pre-fusion fragments (dN/dS vs. pre-fusion sequences, see METHODS). We expect neofunctionalization to be marked by initial diversifying selection followed by purifying selection; however, most fusions identified exhibit signatures of positive selection under both regimes. We note that uncorrected dN/dS is fairly coarse^51^ and the stipulation of an exact 54 base pair match throws out any variation which might be occurring in the very beginning or end of given genes driving down variation which might be occurring. Even with these caveats, the consistent identification of signatures of positive selection seem to indicate many of these deletion-born fusions are still in the process of exploring sequence space for function.

To illustrate one example in more depth, we turn to *narX*, a putative nitrate reductase gene in MTBC genomes. In 182,154 Mycobacterial genomes, *narX* is 1,915 bp long, but in 59 other genomes we identify the prefix and suffix 27-mers separated by 6.6 kb, a 4.6 kb increase from the intact gene (**Figure 4F**). A rooted phylogeny built from sampled genomes shows that they form a monophyletic clade, all belonging to *Mycobacterium canettii*, a smooth tubercle bacillus regarded as one of the closest environmental relatives of transmissible MTBC members.^52^ When we align the *M. tb.* H37Rv *narX* coding sequence to a representative *M. canettii* genome, we find that the “split” *narX* corresponds to a composite of three genes in a nitrate reduction operon and retains three recognizable domains (PF00384: molybdopterin oxidoreductase; PF02613 and PF02665: nitrate reductase delta/gamma subunits). This organization is consistent with a scenario in which a deletion-driven fusion assembled *narX* from a previously distinct nitrate reduction operon. Latent *M. tb.* infections are characterized by long-term survival in hypoxic granulomas, where nitrate respiration supports energy generation in the absence of oxygen.^53–55^ In *M. tb.*, in-vitro experiments demonstrate nitrate reduction is dominated by *narGHJI*, with a minimal role for *narX*.^54^ Although no function has been identified for this fusion, the *narX* example highlights how deletions can rewire core modules and may provide the raw material for the development of novel coding sequences.

The identification of multiple putative deletion-born fusions across the bacterial tree of life indicates the proposed mechanism is shared across bacteria; however, the type strains used were specifically chosen because these species (*E. coli*, *M. tuberculosis*, *N. gonorrhoeae*, *C. jejuni*, *S. pneumoniae*) are some of the most well-studied and consistently sequenced bacteria on the planet. As a result, the genetic variation belonging to these clades is deeply sampled; a reality sadly not shared by most bacteria.^56^ To further clarify the role that sampling bias plays in the ability of the prefix-suffix approach to capture deletion-born fusions across diverse bacterial proteins we turned from a genome-first to a gene-first approach.

### Detection of both deletion-born fusions and broader structural variation is dependent on sampling depth

We next applied the prefix-suffix approach to subsampled collections of 100,000 protein coding ORFs extracted from all 54,630 “complete” RefSeq genomes available at the time of analysis (**Figure 5A**). These genomes contained a total of 219,648,508 protein coding ORFs, these ORFs were clustered into 23,126,961 protein families (see METHODS). Concordant with prior results, we found that most (60.78%) of the protein families have only one sequence within them, so-called “singletons” (**Supplementary Figure 7A**).^57–59^ This observation can partially be explained by the bias of which bacterial genomes are selected for sequencing, assembled with high-confidence, and then annotated. Indeed, we find that the top 20 represented species make up about 35% of the 16,774 species sampled across the dataset (**Supplementary Figure 7B**).

**Figure 5.**
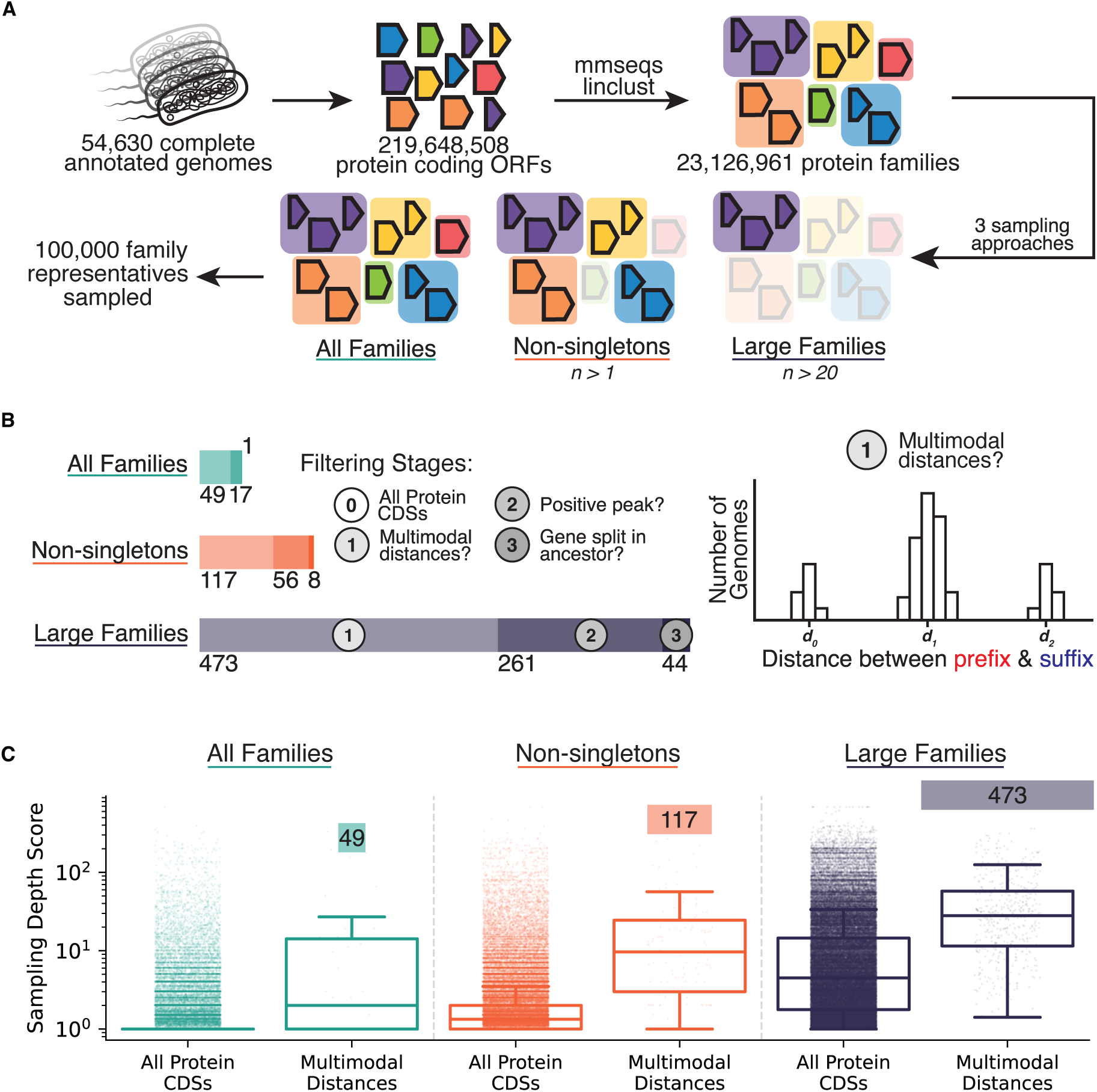
Identification of deletion-born fusions scales with genomic sampling depth. (A) Sampling strategy schematic. Protein-coding ORFs from 54,630 complete RefSeq genomes were clustered into families, and 100,000 families were sampled under three strategies: "All Families" (uniform), "Non-singletons" (≥2 members), and "Large Families" (>20 members). (B) Filtering cascade showing genes passing each stage. "Large Families" yields 44 deletion-born fusions versus 8 ("Non-singletons") and 1 ("All Families"). Cartoon schematic of multimodal distance filtering stage is depicted on the right. (C) Sampling depth score (total genomes / total unique species) for all sampled proteins and multimodal proteins in each sampling strategy.

We sampled 100,000 protein families and selected representatives from each in three distinct ways. In the first, all protein families were given an equal chance of being drawn (“All Families”). In the second, only those families with at least one other sampled member were sampled (“Non-singletons”). In the third, only families with more than 20 sequences in their family were considered (“Large Families”).

Applying the prefix-suffix approach to these 300,000 proteins, we identify increasing deletion-born fusions when moving to more represented protein families (see METHODS). Only 1 gene was identified in “All Families”, 8 genes in “Non-singletons” and 44 genes in “Large Families” (**Figure 5B**, **Supplementary Table 2**). Importantly, these same trends are also observed for the intermediate filtering stages: “Large Families” has not only the most putative deletion-born fusions, but also the most proteins where some structural rearrangement was observed (473 multimodal distances vs. 49 in “All Families”; 217 putative gene insertions vs. 16 in “All Families”).

We sought to examine the extent to which this disparity reflects sampling depth rather than biological distribution by defining a sampling depth score as the total number of genomes containing members of that family divided by the number of unique species represented. A score of 1 indicates a protein family sampled either once or broadly across many species, whereas higher scores indicate repeated sampling within the same species and thus deeper lineage-specific coverage.

Larger protein families were enriched for more deeply sampled species. Protein families in the “Large Families” category exhibited a mean sampling depth score of 15.64, indicating that a typical protein in this set is represented, on average, by 15–16 genomes from the same species (**Figure 5C**). This depth was significantly greater than that of the “Non-singletons” (mean = 3.31) and “All Families” (mean = 1.95) categories (Mann–Whitney U test, p < 10^-30^ for all comparisons), explaining the greater power to detect structural variation in more deeply sampled protein families.

Proteins exhibiting multimodal prefix-suffix distance distributions showed significantly higher sampling depth than the full set of proteins drawn with that strategy (“All Families” mean = 24.15, “Non-singletons” mean = 28.64, “Large Families” mean = 50.53; Mann–Whitney U test, p < 10^-12^ for all comparisons). This pattern indicates that, even controlling for sampling strategy, the detection of structural variation is biased toward protein families with deeper within-species sampling.

Together, these results demonstrate that the apparent enrichment of deletion-born fusions in certain datasets is driven by sampling depth rather than biological prevalence. Detecting these events requires observing both pre-deletion and post-deletion states within the same lineage, which in turn demands not only broad phylogenetic sampling but also dense sampling within species. As genomic databases continue to expand and sampling becomes deeper across the bacterial tree of life, we anticipate that deletion-born fusions will be revealed as more widespread than currently detectable.

## DISCUSSION

In this work, we describe a distinct route to gene birth in which adaptive deletions generate fusion ORFs as by-products. These “deletion-born fusion genes” inherit their initial frequency from a beneficial structural change rather than drifting from rarity and are assembled from pre-existing coding material rather than arising de novo.

This mechanism complements existing models while occupying a distinct niche. Duplication and diversification require maintaining redundant copies in genomes under strong streamlining pressure, overprinting must preserve the ancestral reading frame; and horizontal gene transfer introduces novelty but defers origin to an external donor. Classical gene fusions generally persist only when the fusion is beneficial.^60^ By contrast, deletion-born fusions arise as incidental consequences of selection acting at the level of genome architecture rather than protein function. They require no external material, no long-term maintenance of redundancy, and no immediate benefit of the fused product. Since these fusions are assembled from existing sequences, they are more likely to produce biophysically viable proteins than sequences emerging de novo from non-coding DNA, as proposed for novel genes arising in eukaryotes.^61,62^ Our simulations formalize the hitchhiking advantage showing that elevated starting frequency is the dominant determinant of whether functionalization occurs before loss.

We document this process across multiple evolutionary timescales. In the Lenski LTEE, we observe a large deletion that rapidly sweeps to fixation, generating a novel fusion gene in the process; a convergent deletion at the same locus suggests selection acted on the deletion rather than the gene. At a longer timescale, we identify deletion-born fusions arising during the *Mycobacterium tuberculosis*-*M. bovis* divergence, demonstrating that such fusions can persist across speciation events. Finally, by screening millions of bacterial genomes, we identify putative deletion-born fusions across diverse bacterial clades, indicating that this mechanism reflects a general opportunity for bacterial genome innovation.

We have not, however, identified any clear novel beneficial function to an identified deletion-born fusion. The *yjcO-lysU* fusion in the LTEE shows no nucleotide variation and the *acrR-glcD* and *mlaE-htpX* fusions in the MTBC lack catalytic domains and show minimal variation. Most candidates from our tree-of-life screen exhibit dN/dS ratios consistent with diversifying rather than strong purifying selection, suggesting they remain in the early stages of sequence exploration. The fusions identified here likely represent snapshots of this process, in which proteins persist long enough to sample sequence space but have not yet undergone strong functional refinement. Furthermore, while the genome streamlining literature provides strong indirect evidence that deletions are often beneficial, we have not proven any specific deletion was advantageous and its accompanying fusion was neutral or deleterious.

The absence of clearly functional fusions may reflect biology, but it may also reflect the conservatism of our approach. Our survey provides a lower bound on the prevalence of deletion-born fusions. A single nucleotide mutation in either the prefix or suffix k-mer is sufficient to exclude a genome from the search, so fusions that have accumulated terminal mutations are systematically missed. The problem is compounded by the biology of the events we are seeking deletions often arise through recombination between repetitive sequences, yet repeats are precisely where short-read assemblies tend to break, placing true deletion junctions on contig edges and preventing us from measuring the distance between them.

Future studies employing more sensitive indices could relax the exact-match requirement, potentially revealing functionalized fusions that have since diverged at their termini.

Sampling bias further constrains detection. We show that detection scales with sampling depth: the “Large Families” category, drawn from deeply sampled protein families, yielded 44 deletion-born fusion candidates compared to just 1 in the uniformly sampled “All Families” category. This disparity likely reflects the requirement to observe both pre- and post-deletion states within the same dataset, rather than true biological enrichment in well-studied species. Current genomic databases are dominated by culturable, human-associated pathogens; environmental lineages remain sparsely sampled. This creates a double bind: we both lack the breadth to detect fusions deep in the bacterial tree, and the depth to detect recent fusions in poorly sampled lineages. We expect organisms whose lifestyles predispose them to deletions (i.e. intracellular pathogens) to be enriched for deletion-born fusions, yet many remain too sparsely sequenced to test this hypothesis.

This is not merely a limitation of our approach. Any broad computational analysis of existing databases will inherit these gaps, a fact particularly pertinent given the recent proliferation of protein language models. Our protein family analysis demonstrated that most bacterial proteins in high-quality genome collections remain singletons, solely sampled once. We attribute this to the same sampling bias: we simply have not conducted deep, high-quality sampling across bacterial diversity. Targeted sequencing of underrepresented taxa will be necessary to close these gaps, underscoring that large-scale computational analyses are only as powerful as the underlying biological data relied upon.

The prefix-suffix k-mer approach developed here has utility beyond deletion-born fusions. It enables detection of recurrent structural variation at specific loci across multi-million genome collections neither relying on alignments nor prior knowledge of ancestral states. In this study alone, we characterized internal deletions, repeat prophage insertions, and variable gene cargo in an uncharacterized mobile element. More broadly, the method should be applicable to any process that alters the genomic distance between conserved flanking sequences: novel bacterial or phage introns,^63,64^ integron cassette expansion,^65^ variable plasmid cargo,^66^ or structural variation in complex metagenomic communities.^67^ We have not yet characterized which protein domains or functional categories are enriched among structurally variable loci; such meta-analyses are a natural extension. As sequence databases expand in scale and complexity, alignment-free approaches may become the only tractable means of extracting biological signal from the noisy chorus of bacterial diversity.

Our characterization of deletion-born fusions carries implications beyond evolutionary biology. In clinical microbiology, large deletions are frequently observed during adaptation to host environments;^21,22^ our results suggest that some of these events may generate novel proteins with unpredictable properties. In synthetic biology, rational protein design often proceeds by domain shuffling fusing functional modules from different proteins to create chimeras with new activities.^68^ Understanding how nature assembles and filters such chimeras may inform future efforts to engineer functional novelty.

Our findings recast the deletional bias that pervades bacterial genome evolution as a possible wellspring from which bacterial innovation may arise. By fusing distant sequences into new combinations, deletions may contribute to the vast and still largely uncharacterized diversity of bacterial proteins.

## RESOURCE AVAILABILITY

### Lead contact

Requests for further information and resources should be directed to and will be fulfilled by the lead contact, Arya Kaul.

## Materials availability

This study did not generate new unique reagents.

## Data and code availability

Section 1: Data

- This paper analyzes existing, publicly available data.
- Lenski LTEE

- Metagenomic data downloaded from PRJNA380528
- Clonal sequencing data downloaded from (https://github.com/barricklab/LTEE-Ecoli)
- *Mycobacterium tuberculosis*/*bovis*

- Genomes for *M. tb.* were downloaded from PRJNA719670, PRJNA480888, PRJNA436997 and PRJNA421446.
- Genomes for *M. bovis* were downloaded from PRJNA832544.
- Prefix-Suffix Screen
- AllTheBacteria was downloaded from the OSF repository (https://osf.io/ZXFMY/overview)

Section 2: Code

- Code for analyses besides the prefix-suffix k-mer screen is available at: https://github.com/baymlab/deletion-born-fusion-manuscript
- Code for the prefix-suffix k-mer screen is available at: https://github.com/aryakaul/prefixsuffix-kmer

Section 3: Additional Information

- Any additional information required to reanalyze the data reported in this paper is available from the lead contact upon request.

## ACKNOWLEDGMENTS

The authors would like to warmly thank all past and present members of the Baym and GenScale team for both illuminating conversations and additional scientific insight. This work was supported by NIGMS of the National Institutes of Health (R35GM133700 and R35GM156320), the David and Lucile Packard Foundation, the Pew Charitable Trusts, and the Alfred P. Sloan Foundation. The prefix-suffix approach is based upon research performed in France within the GenScale team at the Inria Center at Rennes University. The work was supported by a Chateaubriand Fellowship of the Office for Science & Technology of the Embassy of France in the United States and a mobility grant from the Collège doctoral de Bretagne. This research was supported by the French National Research Agency (ANR) under Grant ANR-24-CE45-1226 for the REALL project (KB).

## AUTHOR CONTRIBUTIONS

Conceptualization, A.K., K.B., and M.B.; methodology, A.K., K.B., and M.B.; investigation, A.K., F.R., K.B., and M.B.; writing—original draft, A.K. writing—review & editing, A.K., F.R., K.B., and M.B.; funding acquisition, A.K., K.B., and M.B.; resources, K.B. and M.B.; supervision, F.R., K.B., and M.B.

## DECLARATION OF GENERATIVE AI AND AI-ASSISTED TECHNOLOGIES

During the preparation of this work, the author(s) used ChatGPT and Claude to assist with writing code for analysis. After using this tool or service, the author(s) reviewed and edited the content as needed and take(s) full responsibility for the content of the publication.

## SUPPLEMENTARY NOTE(S)

### Estimation of yjcO-lysU/deletion selection coefficient

For a haploid population in which a novel beneficial allele has fitness 1 + *s*, and the wildtype has a fitness of 1, the allele-frequency dynamics are given by:

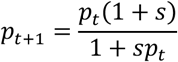

Where *p_t_* is the frequency of the novel allele at generation *t*. This can be approximated to the differential equation, a classical result from Kimura:^69^

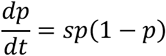

Integrating over this equation and solving for *t* gives:^69^

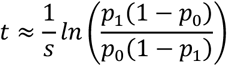

Conditioning on the allele fixing, we can set 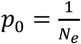 and 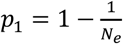. For large effective populations, this yields the commonly used approximation:

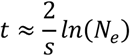

Solving for *s* gives:

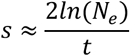

From Good et al.,^39^ we estimate 𝑁_&_ = 10^’^, from our metagenomic analysis, we estimate 𝑡 = 500:

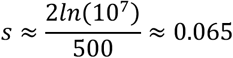

Yielding the estimated selection coefficient of 6.5% for the *yjcO-lysU* fusion/deletion.

## METHODS

### Simulation Framework

We simulated the fate of a novel gene using a minimal forward-time Wright–Fisher model with a fixed haploid population size N = 10^6^ and two initial states: 0 (no fusion), and 1 (non-functional novel gene). State 1 carried a fitness cost *c* (relative fitness 1-*c*), while states 0 had fitness 1. State 2 is when the novel gene functionalizes, and no individual started at this point.

Simulations were initialized with the fusion in state 1 at frequency *p_init_* and the rest at state 0. Populations were updated each generation by fitness-proportionate multinomial sampling (selection + drift). After “reproduction”, individuals in state 1 could functionalize (transition from state 1 to state 2) with probability *p_func_* per generation, and novel gene-bearing individuals in state 1 can lose the gene (transition to state 0) with probability *p_purge_* per generation. Each replicate was run until the novel gene was either lost or functionalized (defined as the first appearance of any state 2 individual, with a maximum of 100,000 generations.

For parameter sweeps (**Supplementary Figure 1**), we evaluated a log-spaced grid (20 points) of *p_func_* from 10^-16^ to 10^-2^ and *p_purge_* from 10^-8^ to 10^-2^ for each combination of *c* and *p_init_* running 100 replicate simulations per grid point with a fixed seed of 42. Simulations were implemented in Python using numpy^70^ and results were saved as matrices of functionalization probabilities. Visualizations provided by seaborn.^71^ Code for simulations is available in the project repository.

### LTEE Data Analysis

We obtained mutation call files for LTEE clonal isolates from the Barrick lab LTEE-Ecoli repository (https://github.com/barricklab/LTEE-Ecoli). For each .gd file, we used gdtools APPLY to generate an isolate-specific genome sequence by applying the curated variants to REL606. We produced per-isolate FASTA outputs. These isolate-specific assemblies were used for consistent gene calling and pangenome construction across all isolates.

To provide standardized gene models for pangenome inference, we annotated each isolate-specific FASTA with Prokka,^72^ producing per-isolate GFF files. These GFFs were used as the input to Panaroo.^73^

For each of the twelve lineages, we ran Panaroo on the set of clonal isolate annotations for that population together with the corresponding ancestral annotation (Anc+ or Anc−). Panaroo was run with --merge_paralogs enabled and with stringent similarity thresholds (e.g., --len_dif_percent 0.98, --threshold 0.98, --family_threshold 0.7), using the “moderate” clean mode.

To identify candidate novel genes arising during evolution, we analyzed each population-specific Panaroo run using a custom script. Briefly, isolate identifiers were mapped to LTEE metadata (population and generation), and Panaroo’s gene_presence_absence.csv and struct_presence_absence.Rtab were scanned to identify gene clusters whose first appearance occurred after the ancestor and increased in presence among later isolates (“appearing genes”).

To distinguish gene families plausibly arising from structural rearrangement from those reflecting annotation noise or near-identical ancestral sequence, we filtered candidate appearing genes by sequence similarity to the REL606 ancestor. For each candidate gene cluster, we extracted its representative nucleotide sequence from pan_genome_reference.fa and aligned it to the ancestral REL606 genome with BLASTN.^74^ We computed the fraction of the candidate gene covered by its single largest BLAST match to REL606 and removed candidates with >85% coverage by the best match, consistent with those being largely ancestral sequence rather than novel junction-derived sequence.

*yjcO-lysU* was identified in this manner and its genomic context from the ancestor was visualized using DNA Features Viewer.^75^ RNA-sequencing and Ribosome profiling were downloaded from Favate et al. and processed according with the same approach as published.^40^ DNA Features Viewer was again used to visualize the reads to the relevant clonal genome.

For candidates that passed the REL606 similarity filter, we extracted a short sequence spanning the putative novel junction. When BLAST produced two distinct hits to REL606, we converted the two hit intervals to BED format and used bedops^76^ with a 30 bp range to identify the local overlap/adjacent junction region; when BLAST produced a single hit, we extracted a 30 bp window around the inferred boundary of the match (depending on hit orientation/endpoint). The corresponding subsequence was extracted from the candidate gene sequence and retained as a “junction” FASTA. To query the relative fraction of the population with a specific variant, both the junction FASTA and the original sequence were aligned with minimap2^77^ to all metagenomic reads from Good et al.^39^ The relative fraction of reads supporting each variant was computed and results were visualized with seaborn.

Domain annotation was performed by using hmmsearch (version 3.4) on the translated protein sequence against Pfam-A database (version 38).^78,79^

To analyze the RNA-seq and Ribo-seq coverage changes around large structural variants, we downloaded a LTEE structural variant table from https://barricklab.org/shiny/LTEE-Ecoli/ on 2025.06.30. We first restricted to the 12 sequenced endpoint clones sequenced in the RNA/Ribo dataset and selected events annotated as deletions or substitutions (DEL or SUB) larger than 1 kb. Because the deletion coordinates were reported in REL606 reference coordinates, we transferred breakpoints into each evolved clone’s coordinate system using whole-genome alignment block coordinates (*.coords, generated by nucmer^80^). For each clone, we parsed alignment blocks describing reference-to-query interval mappings and mapped each breakpoint by locating the corresponding block (allowing a 50 bp tolerance) and applying the within-block offset; events with neither breakpoint mappable were excluded, while events with only one breakpoint mappable were evaluated using a one-sided window around the mapped breakpoint.

Per-base RNA-seq and Ribo-seq coverage was provided as strand-specific depth files for two replicates per clone and for ancestral controls. For each deletion and each window size, we extracted coverage across the mapped interval, averaged coverage across all available replicate and strands to obtain a single mean coverage value and then computed the log_2_ fold change of evolved versus ancestral mean coverage in the same window. To assess spatial scale, we repeated this analysis across multiple window sizes (100 bp to 50 kb). As a matched control, for each clone and window size, we sampled the same number of random genomic windows, excluding regions overlapping deletion windows (including flanks), mapped those windows into clone coordinates using the same alignment-based procedure, and computed coverage log_2_ fold changes identically. Distributions of absolute log_2_ fold change values were then compared between deletion-flanking windows and matched random windows for RNA-seq and Ribo-seq.

### Mycobacterium tuberculosis Complex Analysis

We first downloaded the 36 clinical *M. tb* genomes assembled in Marin et al. 2022 from NCBI using BioProjects PRJNA719670, PRJNA480888, PRJNA436997 and PRJNA421446.^42^ The 10 *M. bovis* genomes used were also downloaded from NCBI using BioProjects PRJNA832544.^43^ Genomes were retrieved in FASTA format and organized per isolate. The M. tuberculosis H37Rv reference genome (accession AL123456) was also downloaded in GenBank format and used as a reference for downstream analyses.

To enable consistent gene-content comparisons across isolates, we predicted protein-coding genes de novo for all genomes using pyrodigal^81^ and exported predicted proteins as amino acid FASTA files. All predicted proteins across genomes were then clustered into gene families using MMseqs2.^82^ We created a single MMseqs2 sequence database from all proteins, clustered sequences using identity-based clustering (minimum sequence identity 0.7), and exported cluster assignments as a tab-delimited table. These clusters were used to define gene families and identify accessory families whose presence varied across *M. tb* and *M. bovis* isolates.

To prioritize candidate deletion-born fusions, we examined accessory gene families with lineage-restricted presence patterns and then performed nucleotide-level mapping in genomes lacking the gene family. For each candidate family, we extracted representative nucleotide sequences and aligned them to the corresponding genome FASTA files using BLASTN. BLAST hits were converted to genomic intervals, merged to identify discrete matching regions, and filtered to enrich for signatures consistent with deletion-born fusion rather than simple absence, fragmentation, or duplication. Specifically, retained candidates required at least 80% total aligned coverage across merged hits, a best single-hit length not exceeding 80% of the gene length, and multiple hits separated by at least 1 kb in the target genome. Candidates passing these criteria were treated as putative deletion-born fusion genes. Domain analysis was done as before for the *yjcO-lysU* fusion.

To place candidate events in an evolutionary context, we constructed a core-genome alignment and phylogeny using Parsnp^83^ with H37Rv as the reference, using the set of *M. tb* and *M. bovis* assemblies analyzed above. The resulting phylogeny was visualized in iTOL^84^ and BLAST results were visualized using DNA Features Viewer.^75^

### Prefix-Suffix K-mer Screen

We extracted prefix and suffix k-mers from each query gene using coding nucleotide FASTA inputs. For each gene sequence, we took a k-mer of length k (k = 27) from near the 5′ end and a second k-mer of length k from near the 3′ end, excluding a small buffer region from each terminus to avoid start and stop codons and to shift the k-mers out of frame relative to the annotated coding sequence. Specifically, we used a fixed gap distance g = 4 bp: the prefix k-mer was taken from positions g to g+k (4 to 31), and the suffix k-mer from positions −(g+k) to −g.

We queried these k-mers against the AllTheBacteria collection of 2.4 million bacterial isolate assemblies. Assemblies were processed in batches according to their Miniphy’d^85^ output: genome FASTA files were stored in compressed tar.xz archives, and for each archive we concatenated subsets of 500 genomes into temporary multi-FASTA files and built BWA indices on these batches. Indexing was performed with bwa index, producing FM-indices for each genome batch.

Exact k-mer placements were then obtained using bwa fastmap, run with the query k-mer FASTA and a matching k-mer length equal to k (the same value used in extraction). We used a large maximum hit window (-w 99999) to retain all exact match locations reported by fastmap. Fastmap outputs were gzip-compressed and parsed to recover, for each genome, all contig-level match positions for both prefix and suffix k-mers.

For each genome and each query gene, we computed a prefix-suffix distance only when at least one prefix match and one suffix match occurred on the same contig. Distances were computed from the genomic coordinates of the matched k-mers on that contig. Matches split across contigs were ignored, and genomes lacking a same-contig prefix-suffix pair were treated as missing for that gene.

### Multimodal Distance Detection and Clustering

We identified structurally variable genes by clustering the per-genome prefix-suffix distances for each query gene and testing whether the resulting distance distribution was multimodal. For each gene, we aggregated the set of observed “Difference” values across genomes (the prefix-suffix distance expressed relative to the expected gene length) along with their multiplicities, and clustered these one-dimensional values using a density-based algorithm (DBSCAN1D).^86^ Clustering was run with epsilon = 800 (bp) and min_samples = 25, and we treated DBSCAN noise points as outliers (removed from downstream summaries). A gene was called “multimodal” only if DBSCAN identified at least two non-noise clusters, corresponding to two tight peaks in the distance distribution.

To enrich specifically for candidates consistent with deletion-born fusion genes, we applied additional filters to multimodal loci. We required one cluster centered near 0 (the intact allele, where the observed prefix-suffix distance matches the reference gene length) and at least one additional cluster centered at a positive distance greater than 800 bp, consistent with a split ancestral state in which the prefix and suffix k-mers are separated by a substantial intervening segment. Genes with only small positive shifts (e.g., internal deletions) or without an intact-like cluster near 0 were excluded at this stage. The set of genes passing these clustering and distance-peak criteria was carried forward for phylogenetic filtering and downstream analyses.

### Phylogenetic Reconstruction and Ancestral State Inference

To distinguish deletion-born fusions from gene disruption by insertion (**Figure 4B**, filter 3), we performed phylogenetic reconstruction and ancestral state inference using the DBSCAN cluster assignments from the prefix-suffix distance analysis. For each candidate gene passing the multimodality and peak-shape filters, we downsampled genomes to obtain a tractable but representative set for tree building by sampling equal numbers of genomes from each distance cluster (300 total genomes per gene, split evenly across clusters). Sampling was restricted to a high-quality genome set from the ATB to reduce artifacts from fragmented assemblies. For each sampled genome, we extracted the identified prefix and suffix k-mer, as well as the entire interleaving sequence to a new FASTA file and masked that same region in the whole genome sequence.

Trees were constructed from masked sequences using attotree^87^ (an optimized version of Mashtree^88^) with default options on the masked whole genome sequences. To enable rooting, we also added two outgroup genomes per focal species, chosen as close relatives outside the focal clade: for *E. coli* K-12, *Klebsiella pneumoniae* MGH78578 (GCF_000016305.1) and *Salmonella enterica* serovar Typhimurium LT2 (GCF_000006945.2); for *M. tb.* H37Rv, *M. canettii* (NC_015848.1) and *M. caprae* (CP016401.1); for *N. gonorrhoeae* FA1090, *N. lactamica* (NC_014752.1) and *N. meningitidis* MC58 (NC_003112.2); for *C. jejuni* NCTC1168, *C. coli* (NC_022660.1) and *C. lari* (NC_012039.1); and for *S. pneumoniae* TIGR4, *S. mitis* (FN568063.1) and *S. oralis* (FR720602.1). For each of the two possible outgroups included, we identified the genome with the largest mean distance between it and the other leaves and chose that as the outgroup to root the resulting tree at.

For each candidate gene passing the distance-based filters, we inferred whether the ancestral state was “split” or “intact” using a tree-based parsimony approach. Cluster labels were treated as discrete character states, and we reconstructed internal node states by multi-state Fitch parsimony.^89^ The ancestral state for each gene was defined as the parsimony assignment at the most recent common ancestor (MRCA) of the ingroup genomes. Genes were classified as consistent with deletion-born fusions when the MRCA state corresponded to a “split” cluster (the cluster with a positive prefix-suffix difference) and at least one descendant clade carried an “intact” cluster (the cluster with mean difference near 0). Conversely, genes whose MRCA state was “intact” were interpreted as cases where the intact gene was ancestral, and the positive-distance cluster reflects disruption (often by insertion) and were excluded from the deletion-born fusion set.

### Mobile Genetic Element and Prophage Detection

To assess whether structurally variable loci were dominated by insertions of mobile genetic elements (MGEs) or prophages (**Figure 4C**), we aligned the sequence between the matched prefix and suffix k-mers to curated MGE and prophage databases. The MGE database was taken from MGEdb^90^ and the prophage database from Prophage-DB^91^ (bacterial host prophages); the two FASTA sets were concatenated into a single nucleotide BLAST database. For each genome sampled, we extracted the intervening sequence between the matched prefix and suffix k-mers on the same contig (i.e., the sequence whose length drives the positive prefix-suffix distance signal) and queried it against the combined database using BLASTN. For each intervening sequence, we merged overlapping BLAST hit intervals along the query and computed the total number of query bases covered by any hit; the fraction of the intervening sequence explained by MGEs/prophages was then calculated as covered bases divided by query length. These per-genome fractions were summarized by locus and compared between loci whose inferred MRCA state was “split” versus “intact” using a two-sided Mann–Whitney U test.

### Selection Analysis

For the selection analysis on *mlaE-htpX* and *acrR-glcD*, genomes with the intact ORF were identified based on their membership to the cluster with a relative prefix-suffix “Distance” of 0. The prefix-suffix k-mer and intervening nucleotides were extracted, deduplicated, and codon-aligned using MACSE with default options.^92^ A phylogeny of the masked whole genome sequences was built using Parsnp and selection analysis was performed with FUBAR, implemented in the HyPhy package with default options.^93^

We assessed signatures of selection on candidate deletion-born fusions using two complementary dN/dS-style comparisons (**Figure 4E**). First, to estimate selection acting on the observed “intact” allele, we focused on genomes assigned to the cluster closest to zero difference (the intact-like cluster). For each genome we used the extracted locus sequence and deduplicated identical sequences by hashing, retaining a count of how many genomes shared each unique sequence. Each unique sequence was then codon-aligned to the canonical query CDS using MACSE,^92^ and we counted synonymous and nonsynonymous differences across aligned codons, ignoring codons overlapping gaps or ambiguous bases and tracking premature stop gains separately. Per-cluster estimates were computed by summing synonymous and nonsynonymous counts across unique sequences and weighting each sequence by the number of genomes in which it occurred; dN/dS was then calculated as weighted nonsynonymous divided by weighted synonymous changes.

Second, to estimate the degree of divergence expected from the pre-deletion (“split”) state, we constructed “surrogate” genes for genomes assigned to the inferred ancestral cluster (the MRCA cluster from parsimony). For each genome we used the extracted intervening locus sequence and flanking sequence, deduplicated both sequence sets, and mapped each unique removed-region sequence into its parent flank sequence by exact substring matching (allowing reverse complement). We then aligned the canonical query CDS to each unique removed-region sequence using BLASTN and selected a set of high-scoring segment pairs that approximately tiled the query. Each BLAST segment was projected back onto the parent flank coordinates and intersected with ORFs predicted on the flank sequence using pyrodigal; for each segment we selected the ORF with the greatest overlap and stitched the resulting ORF nucleotide sequences together in query order, joining adjacent blocks with “NNN” to preserve codon-phase ambiguity. These stitched constructs were treated as surrogate pre-deletion sequences and were codon-aligned to the canonical query CDS again using MACSE, after which synonymous and nonsynonymous differences were counted as above and aggregated into a weighted dN/dS estimate.

### Protein Family Clustering and Sampling

We constructed a large-scale bacterial protein family catalogue from complete RefSeq genomes to quantify how sampling depth affects detection of deletion-born fusion genes (**Figure 5**). All complete bacterial genomes available in RefSeq at the time of analysis (54,630 assemblies) were downloaded using ncbi-genome-download in GenBank format. From each genome, we extracted all annotated protein-coding sequences by parsing the GenBank feature tables. For each CDS, we extracted both the nucleotide sequence and the corresponding protein sequence. When a translation was provided in the GenBank record it was used directly; otherwise, the nucleotide sequence was translated in-frame. Protein sequences were written to per-genome FASTA files and then concatenated into a single combined protein FASTA for clustering.

Protein sequences were clustered into gene families using MMseqs2 Linclust.^82^ Clustering was performed with a minimum pairwise sequence identity of 80%, a minimum target coverage of 80% (coverage mode 1), and 80 k-mers per sequence, using mmseqs easy-linclust. This procedure yielded 23,126,961 protein families, spanning both singleton and multi-member families. The resulting cluster table and representative sequences were used for downstream sampling and analysis.

To evaluate how database sampling affects detection of structural variation, we defined three complementary protein-family sampling strategies. In the “All Families” strategy, we sampled protein families uniformly at random from the full set of MMseqs2 clusters, including singletons. In the “Non-singletons” strategy, we excluded singleton families and sampled uniformly from families containing at least two members. In the “Large Families” strategy, we restricted sampling to families with more than 20 members, enriching for deeply sampled lineages. For each strategy, we analyzed an equal number of protein families (100,000).

For each protein family, we quantified sampling depth using a sampling depth score defined as the total number of genomes containing members of that family divided by the number of unique species represented among those genomes. A score near 1 indicates a family sampled broadly but shallowly across species, whereas higher values indicate repeated sampling of the same species (e.g., many isolates of a single lineage). Sampling depth scores were used to compare the effective detectability of deletion-born fusions across the three sampling strategies.

---

**Supplementary Figure 1.**
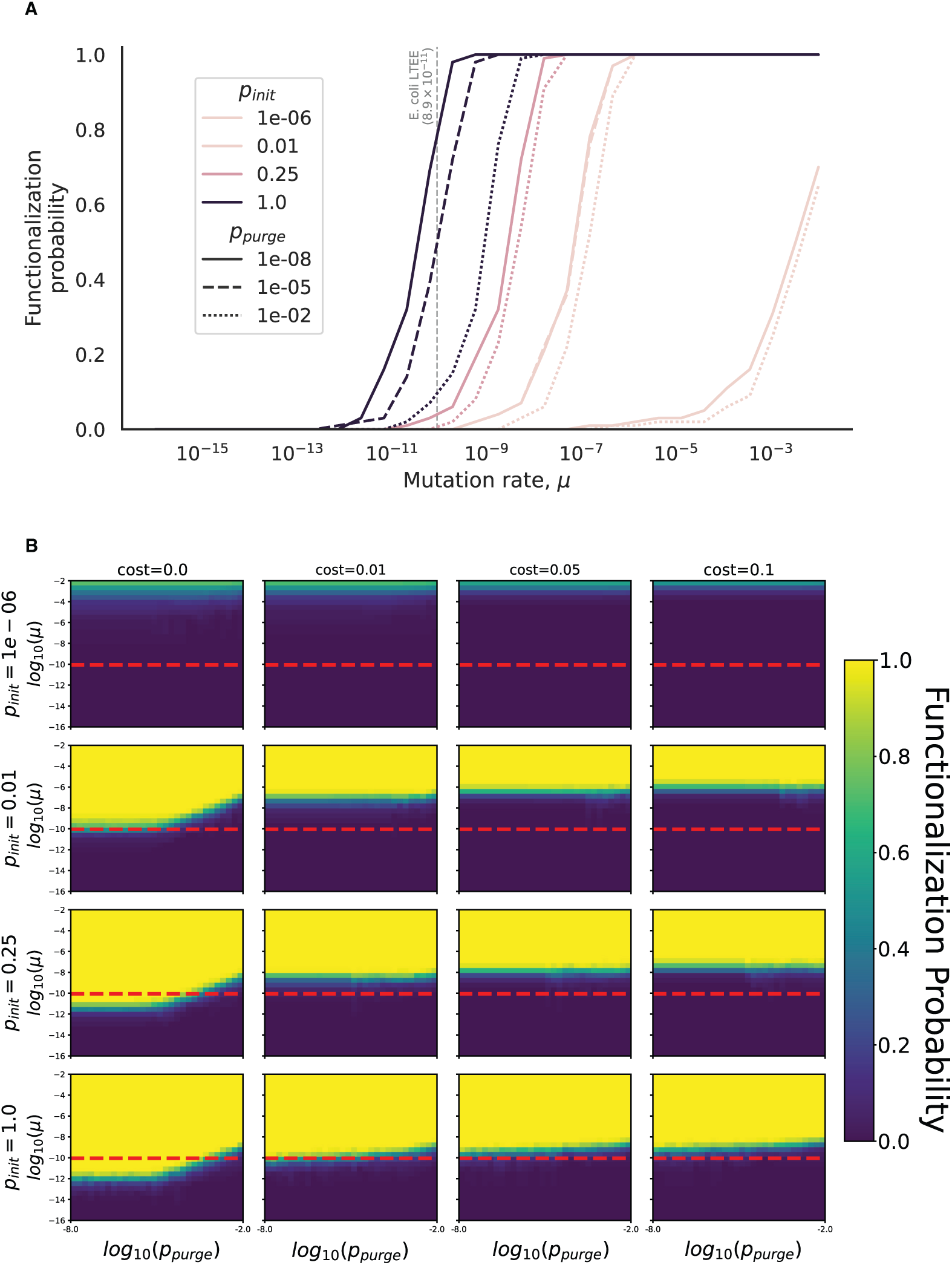
Forward simulations quantify the hitch-hiking advantage (A) Probability that at least one lineage functionalizes as a function of the mutation rate *µ*; colors denote four starting frequencies of the nascent fusion (*p_init_*) and line styles three purge probabilities (*p_purge_*). The dashed vertical line marks the LTEE point-mutation rate. All curves are shown for a fusion gene with a pre-functionalized fitness cost of 0.01. (B) Heat-maps show the same probability across grids of *p_purge_* (x-axis, log10 scale) and fitness cost of the unfixed fusion (*c*, four columns) for the four *p_init_* values (rows); the color of the cell indicates the proportion of times the fusion functionalized before being purged. The dashed red line denotes the LTEE point-mutation rate. Each cell summarizes 100 Wright–Fisher runs of 10^6^ haploids.

**Supplementary Figure 2.**
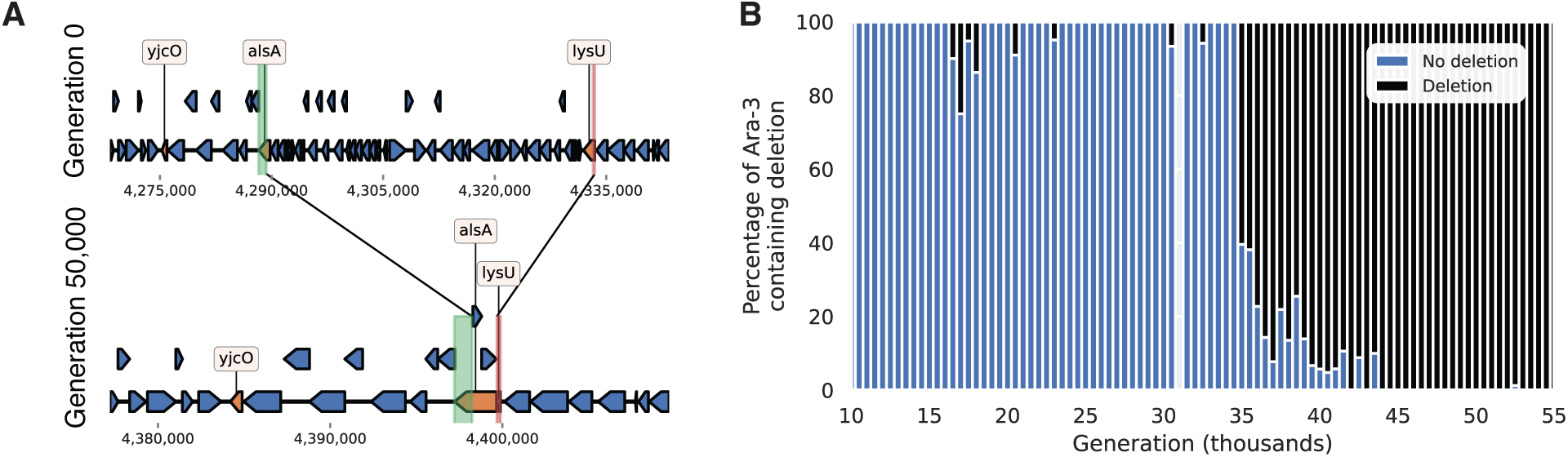
A convergent 43.4 kb deletion sweeps at the same locus in Ara-3. (A) Schematic of the deletion in Ara-3, orange genes highlight the resulting prior genes involved in the deletion. Green and red bars correspond to BLAST alignments to this region.(B) Metagenomic sequencing results showing the fraction of reads supporting the deletion or not.

**Supplementary Figure 3.**
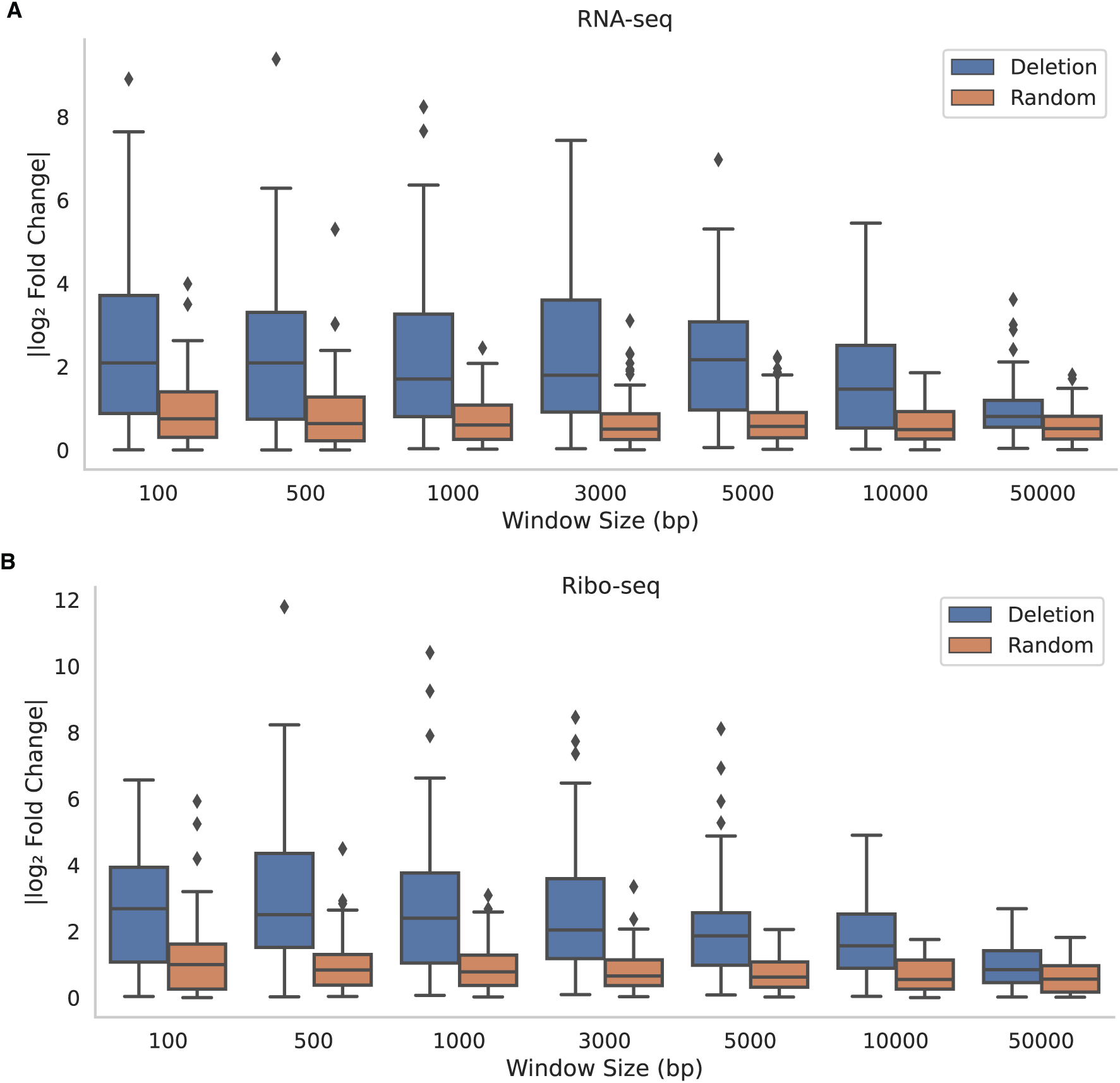
Large LTEE deletions are associated with significant changes to local transcription and translation. (A) RNA-seq log2 fold changes for windows of varying size flanking ≥1 kb deletions (blue) and for randomly sampled windows (orange). Fold changes are calculated between the ancestral strain and the evolved population at generation 50,000. Data downloaded from Favate et al. 2022.^40^ (B) As in (A) but analyzing Ribo-seq data.

**Supplementary Figure 4.**
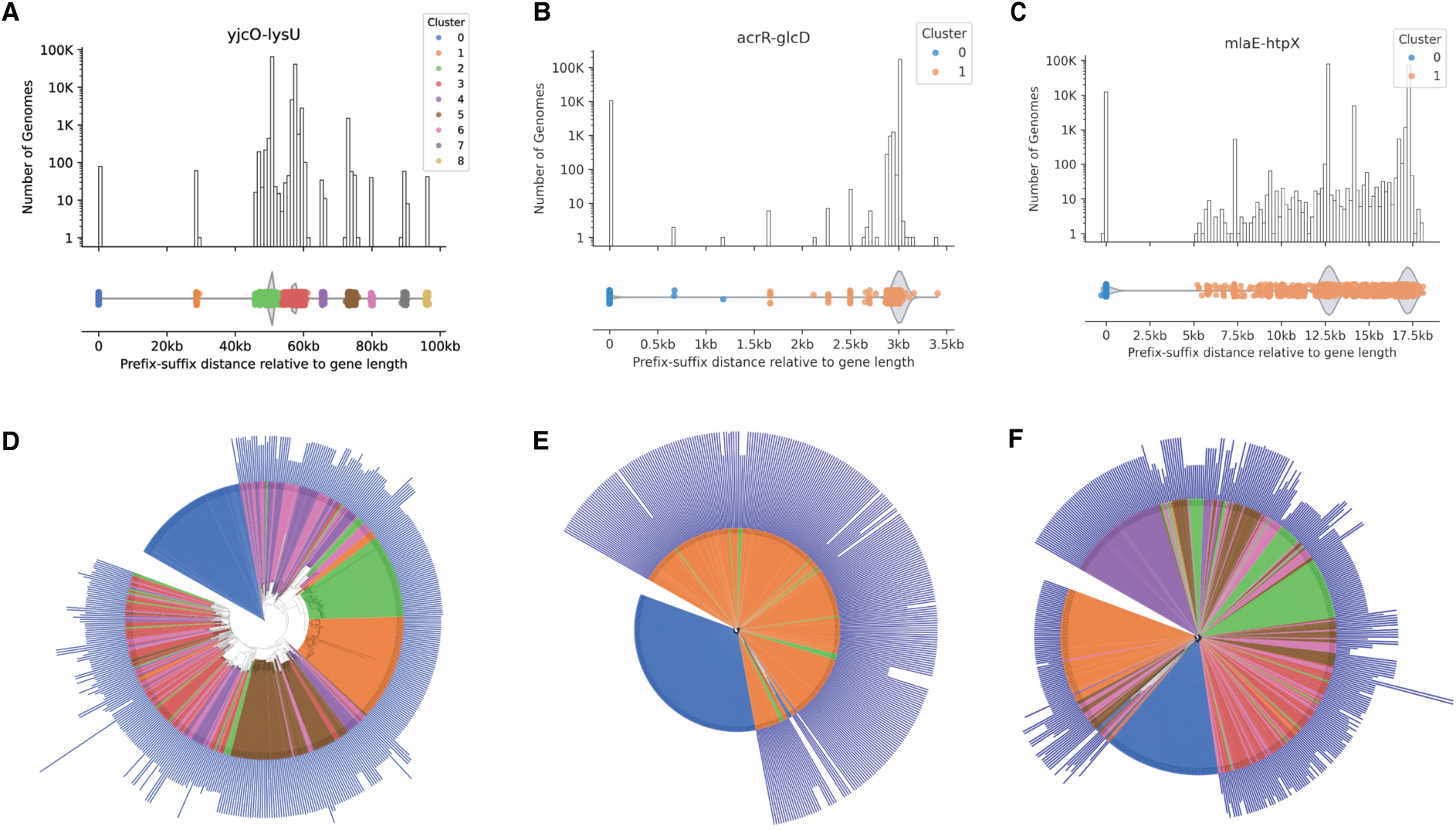
The prefix-suffix approach captures previously identified deletion-born fusions and reveals additional structural variation. (A) Top: Log-scaled histogram of prefix-suffix distances relative to the length of the original *yjcO-lysU* fusion. Bottom: underlying point distribution for histogram. Every point represents a single genome in the ATB dataset; colors correspond to DBSCAN-derived clusters. (B) (**C**) are the same as in (A) but with *acrR-glcD* and *mlaE-htpX* respectively. (D) Distance-metric based phylogeny of 300 randomly sampled genomes with equal genomes sampled across the number of clusters identified. Clades are colored by cluster membership, and blue vertical bars off leaves represent the distance between the prefix-suffix found in that genome relative to the *yjcO-lysU* gene. (E), (**F**) are the same as in (D) but with *acrR-glcD* and *mlaE-htpX* respectively.

**Supplementary Figure 5.**
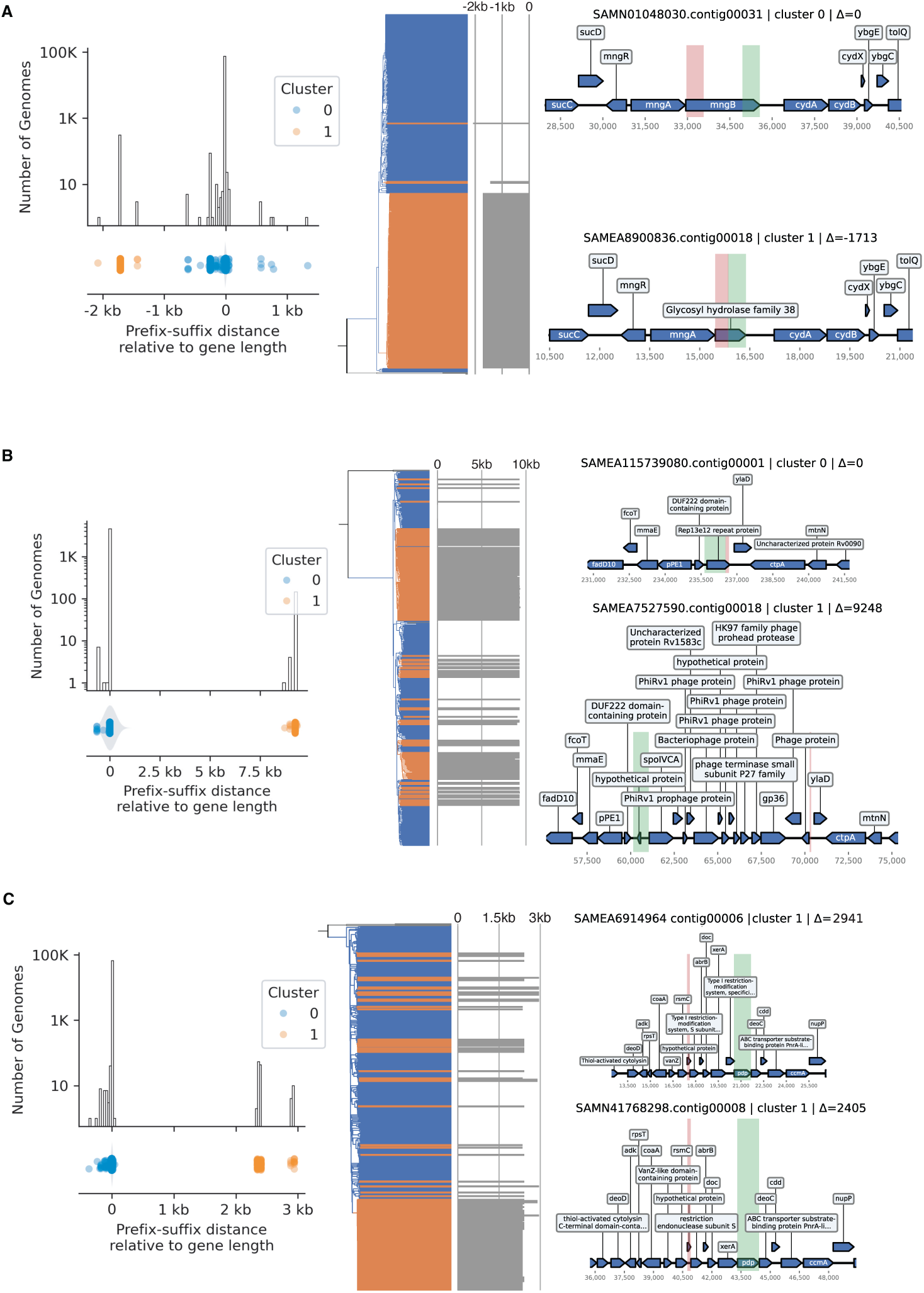
Prefix-suffix approach captures diverse structural variation beyond deletion-born fusions. Each row shows: prefix-suffix distance distribution (left), rooted phylogeny of 300 sampled genomes (middle), and representative genomic contexts (right). (A) Internal deletion in *mngB* (*E. coli* K12). (B) Repeat prophage insertions in *rep13e12* (*M. tuberculosis* H37Rv). (C) Variable gene cargo disrupting *pdp* (*S. pneumoniae* TIGR4); the two Cluster 1 representatives differ by ∼500 bp due to the presence/absence of a Type I restriction system protein downstream of *xerA*.

**Supplementary Figure 6.**
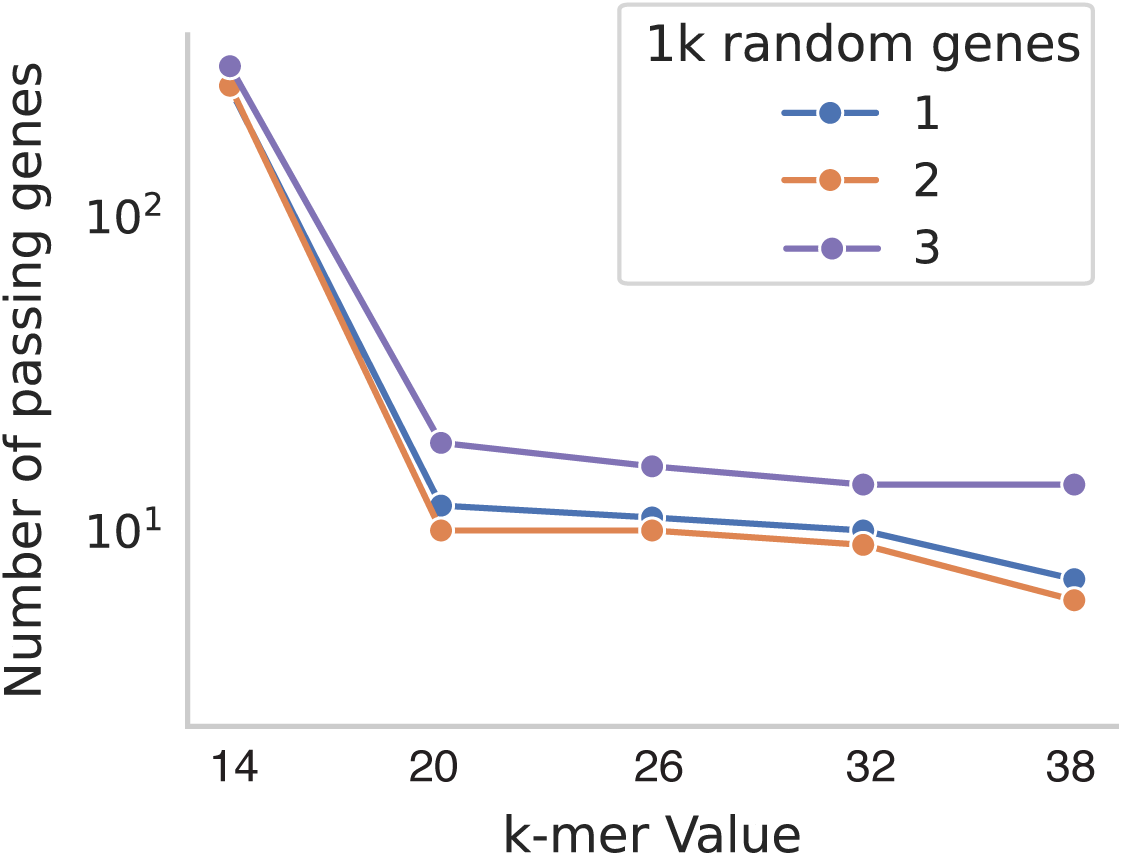
Prefix-suffix approach is robust to values of k ≥ 20. Number of genes with multimodal prefix-suffix distances detected across three random samples of 1,000 RefSeq genes at varying k-mer lengths.

**Supplementary Figure 7.**
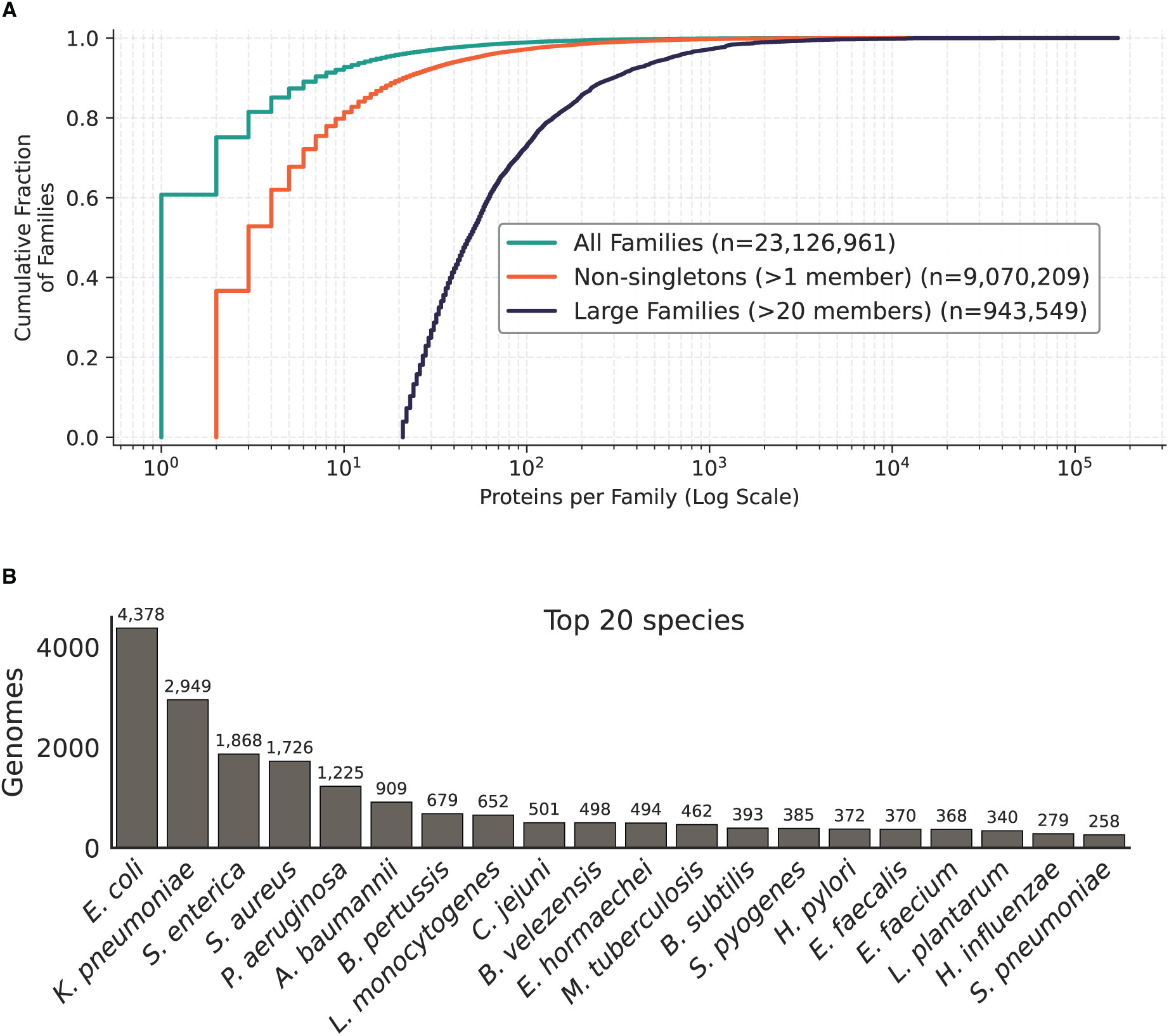
Representation of both protein families and bacterial species are highly skewed in RefSeq complete genomes. (A) Cumulative distribution function of the number of protein members per family. Most represented proteins are singletons (B) Top 20 species represented in the 54,630 complete genomes proteins were pulled from. Count of each is displayed above each bar.

